# Common patterns of gene regulation associated with Cesarean section and the development of islet autoimmunity – indications of immune cell activation

**DOI:** 10.1101/167676

**Authors:** M. Laimighofer, R. Lickert, R. Fürst, F. J. Theis, C. Winkler, E. Bonifacio, A.-G. Ziegler, J. Krumsiek

## Abstract

**Background:** Birth by Cesarean section increases the risk of developing type 1 diabetes later in life; however, the underlying molecular mechanisms of this effect remain unclear. We aimed to elucidate common regulatory processes observed after Cesarean section and the development of islet autoimmunity, which precedes type 1 diabetes, by investigating the transcriptome of blood cells in the developing immune system.

**Methods:** We analyzed gene expression of peripheral blood mononuclear cells taken at several time points from children with increased familial and genetic risk for type 1 diabetes (n = 109). We investigated effects of Cesarean section on gene expression profiles of children in the first year of life using a generalized additive mixed model to account for the longitudinal data structure. To investigate the effect of islet autoimmunity, we compared gene expression differences between children after initiation of islet autoimmunity and age-matched children who did not develop islet autoantibodies. Finally, we compared both results to identify common regulatory patterns of Cesarean section and islet autoimmunity at the gene expression level.

**Results:** We identified two differentially expressed pathways in children born by Cesarean section: the pentose phosphate pathway and pyrimidine metabolism, both involved in nucleotide synthesis and cell proliferation. Islet autoantibody analysis revealed multiple differentially expressed pathways generally involved in immune processes, including both of the above-mentioned nucleotide synthesis pathways. Comparison of global gene expression signatures showed that transcriptomic changes were systematically and significantly correlated between Cesarean section and islet autoimmunity. In addition, signatures of both Cesarean section and islet autoimmunity correlated with transcriptional changes observed during activation of isolated CD4+ T lymphocytes.

**Conclusions:** We identified coherent gene expression signatures for Cesarean section, an early risk factor for type 1 diabetes, and islet autoantibodies positivity, an obligatory stage of autoimmune response prior to the development of type 1 diabetes. Both transcriptional signatures were correlated with changes in gene expression during the activation of CD4+ T lymphocytes, reflecting common molecular changes in immune cell activation.

## Background

Type 1 diabetes is an autoimmune disease in which immune cells destroy insulin-producing beta cells in the pancreas. Loss of beta-cell mass leads to insulin deficiency, impaired glucose tolerance, and ultimately the onset of type 1 diabetes [1]. This process is marked by the appearance of islet autoantibodies to beta-cell-related antigens [2]. Autoantibody seroconversion usually occurs during childhood and early adolescence, with a peak conversion rate between 9 months and two years [3]. Notably, an increasing incidence of type 1 diabetes has been observed within the last few decades, especially in children [4]. Several previous studies indicated transcriptional changes in immune cells caused by the development of islet autoantibodies [5, 6], indicating substantial molecular changes long before the onset of type 1 diabetes.

Children born by Cesarean section have an odds ratio of 1.23 for the development of type 1 diabetes compared to vaginally delivered children [7]. Moreover, Cesarean section has been reported to be associated with a faster progression to type 1 diabetes, but not with an increase the risk for islet autoimmunity [8]. The underlying mechanisms of these associations are not yet fully understood. Some reports indicate that the mode of delivery affects colonization of microbiota in the intestinal tract [9], which in turn affects the developing immune system of infants [10]. Such differences in the gut microbiome and its interaction with the immune system may lead to increased risk of asthma, childhood allergies [11], and autoimmune diseases, such as type 1 diabetes.

Here we investigated the impact of Cesarean section and islet autoimmunity on the immune system by analyzing gene expression data in peripheral blood mononuclear cells (PBMCs) as a readout (Figure 1A). The overall goal of the analysis was to elucidate common immune modulation patterns between an early risk factor, Cesarean section, and the development of autoimmunity. We first investigated the effects of Cesarean section on gene expression profiles of children in the first year of life (Figure 1B). To this end, we analyzed data from several time points in this early period in children with an increased familial risk for type 1 diabetes. A generalized additive mixed model was used to account for the longitudinal information when extracting gene expression differences between children born by Cesarean section and vaginal delivery (Figure 1C). In a second analysis, we compared gene expression levels of children with increased risk of familial type 1 diabetes after initiation of islet autoimmunity with age-matched children who did not develop islet autoantibodies. We then combined both results to identify genes and pathways that were differentially regulated in both analyses, to link Cesarean section and islet autoimmunity at the level of gene expression. Finally, we compared the patterns identified for Cesarean section and islet autoantibodies with gene expression data from activated human CD4+ T lymphocytes to identify candidates for common molecular mechanisms of immune activation.

**Figure 1:**
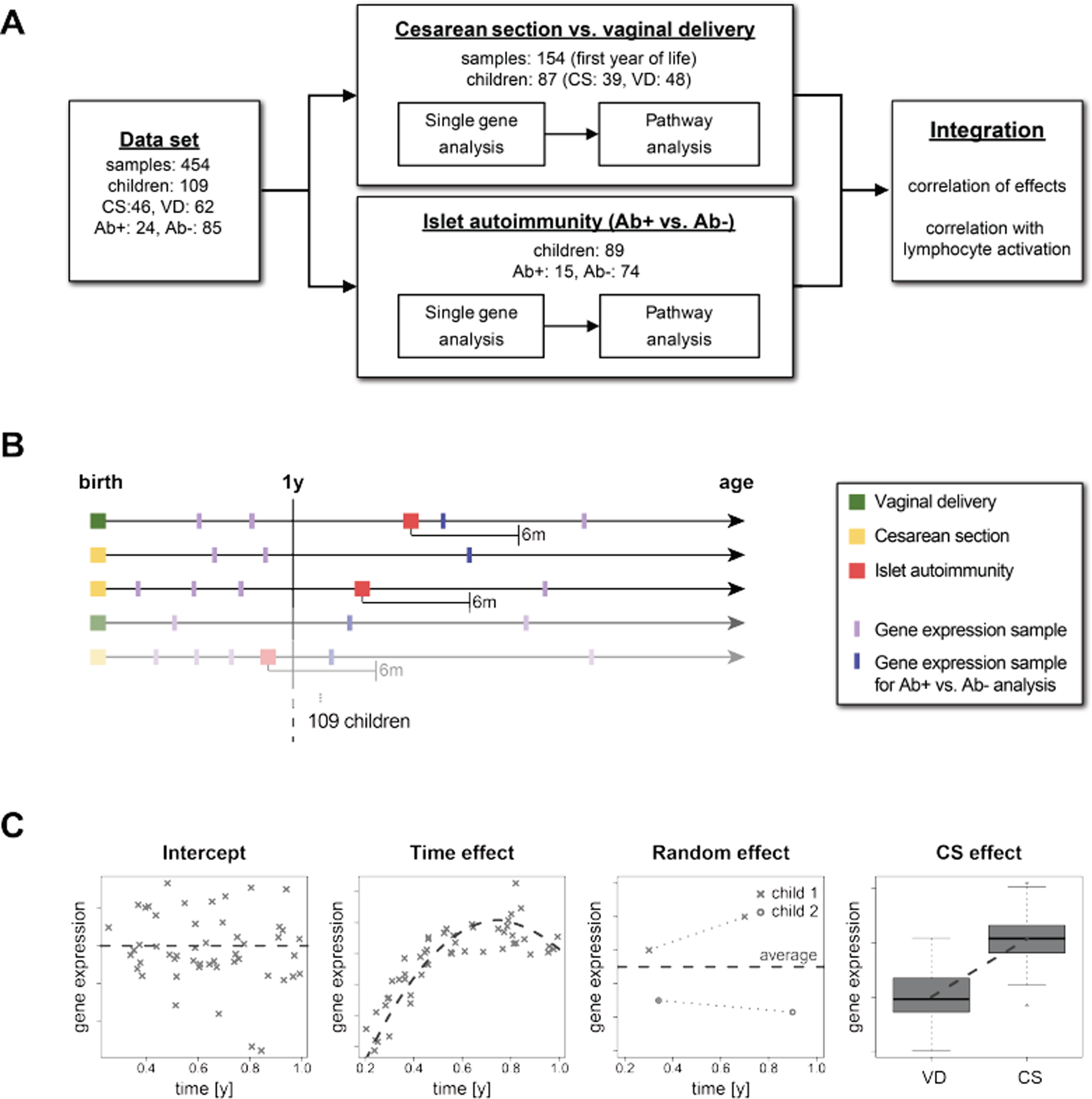
Overview. **A:** Study workflow: Parallel analyses were performed to detect differential gene expression and pathway enrichment for Cesarean section and islet autoimmunity. The results were then compared in a combined analysis and related to expression patterns of lymphocyte activation. **B:** Schematic overview of the longitudinal study design for Cesarean section and islet autoimmunity analyses. **C**: Schematic illustration of the generalized additive mixed effect model (GAMM) to analyze the longitudinal dataset, including intercept, a time effect, a random effect for multiple measurements, and the investigated Cesarean section vs. vaginal delivery effect. Abbreviations: CS = Cesarean section, VD = Vaginal delivery, Ab+ = Islet autoantibody-positive, Ab− = Islet autoantibody-negative.

## Results

### Effects of Cesarean section on the transcriptome in the first year of life

To focus on early postpartum effects on genetic regulation after delivery by Cesarean section, we restricted our analysis to samples from children in their first year of life (Supplemental file 1: Figure S1). This resulted in a total of 154 PBMC gene expression samples (71 Cesarean section, 83 vaginal delivery) from 87 children (39 Cesarean section, 48 vaginal delivery), Figure 1A+B. The number of samples per child ranged between 1 and 4 (Supplemental file 1: Figure S1). Details on the dataset and preprocessing steps are provided in the Methods section.

We used a generalized additive mixed effect model (GAMM) to account for the longitudinal nature of the dataset. The model consisted of three major parts (Figure 1C): (1) the change of expression profiles over time was modeled using a spline-based function, (2) the overall trend of expression values per child was captured by a random mixed effect, and (3) a term representing the Cesarean section effect extracted the association in which we were primarily interested. Applying this model to the data set, we observed an enrichment of low p-values before adjusting for multiple testing (Figure 2A and Supplemental file 2). However, after multiple testing correction by controlling the false discovery rate (FDR) at 0.1, no significant single genes could be identified in this first step (Figure 2B).

**Figure 2:**
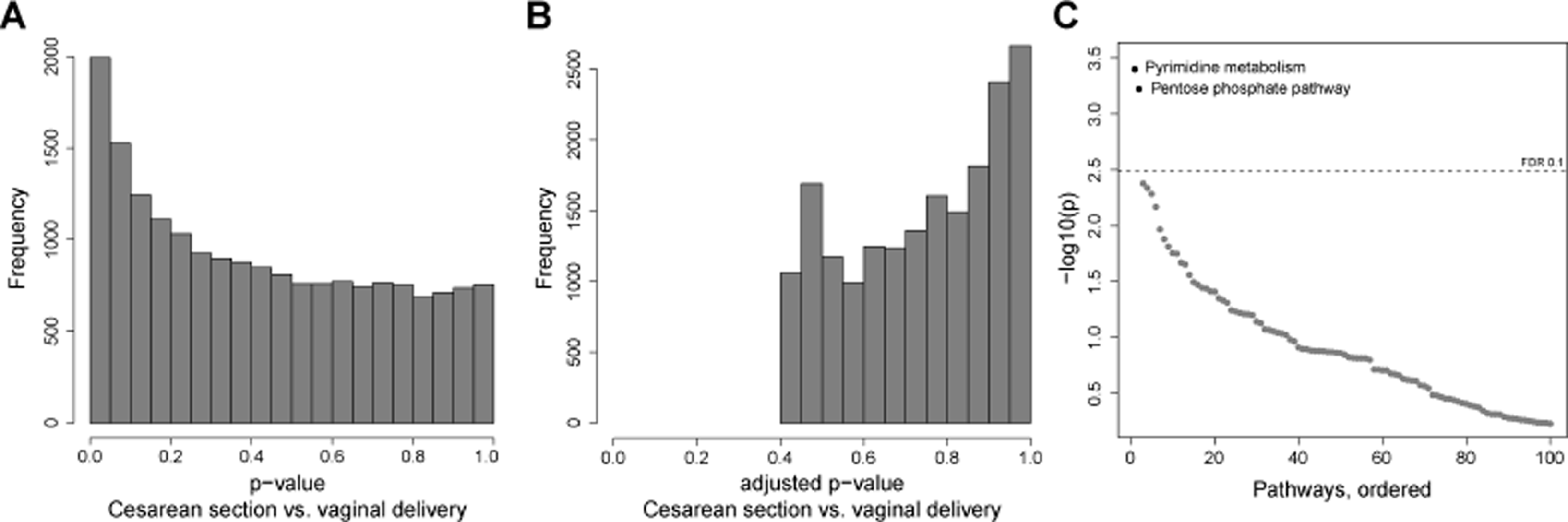
Cesarean section analysis. **A**: Histogram of unadjusted p-values for CS association per gene. **B**: Histogram of p-values after multiple testing adjustment by controlling the false discovery rate. **C:** Sorted −log10(p-values) of pathway enrichment. Dashed line indicates the multiple testing threshold at an FDR of 0.1. Abbreviations: CS = Cesarean section, VD = Vaginal delivery.

To improve statistical power and facilitate biological interpretability, we performed a pathway enrichment analysis based on the KEGG pathway database [12]. From this analysis, we obtained two significantly differentially expressed pathways after multiple testing correction (FDR <= 0.1): “Pentose phosphate pathway” and “Pyrimidine metabolism” (Figure 2C and Supplemental file 2). The “Pentose phosphate pathway” included 27 genes (20 up- and 7 down-regulated genes in children born by Cesarean section) and the “Pyrimidine metabolism” pathway contained 92 genes (57 up-regulated and 35 down-regulated).

Both pathways are well known to be involved in the synthesis of nucleotides during cell proliferation [13, 14]. This indicated that there is a change in expression profiles of proliferation-related nucleotide synthesis pathways as an early consequence of delivery by Cesarean section.

### Effects of islet autoimmunity on the transcriptome

To identify gene expression signatures associated with seroconversion, we compared the earliest sample of children after the development of islet autoantibodies (up to 6 months post-seroconversion; 15 children) with all available age-matched samples of children who did not develop islet autoantibodies (74 children). We applied linear regression to explain gene expression differences induced by islet autoantibody status, corrected for age (see Methods). We detected a strong accumulation of differentially expressed genes (Figure 3A), of which 3,867 were significant after multiple testing correction (FDR ≤ 0.1, Figure 3B). The majority of these genes were found to be up-regulated in children with islet autoimmunity (64% up-regulated vs. 36% down-regulated, Supplemental file 2).

**Figure 3:**
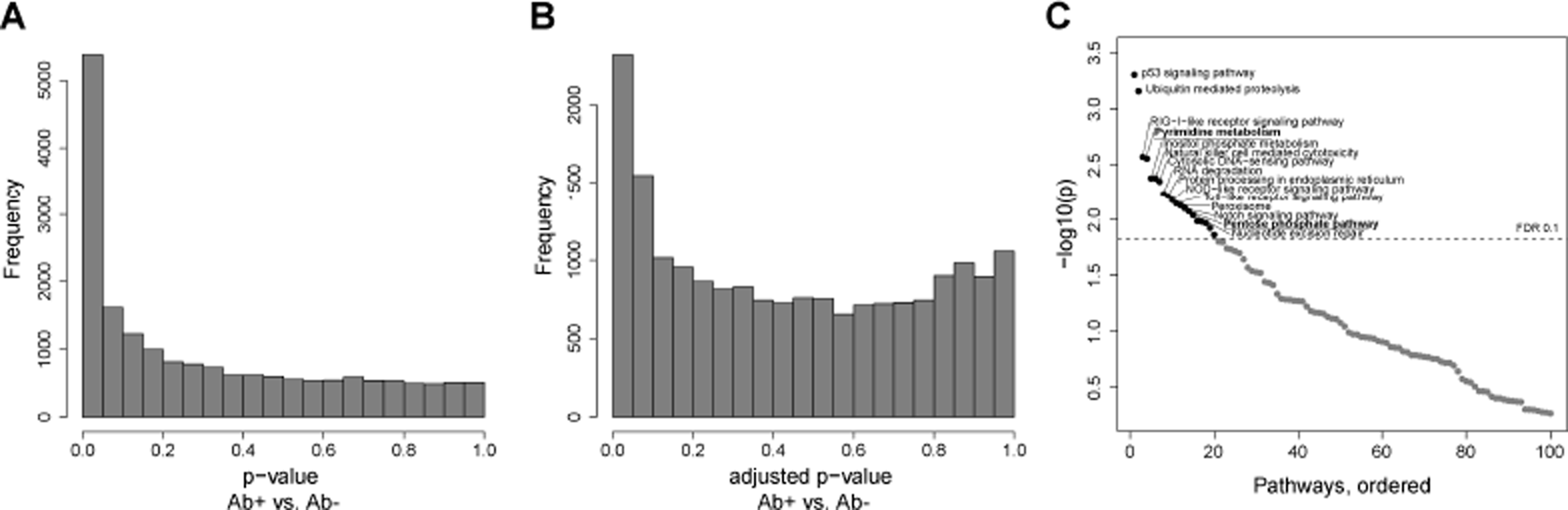
Ab+ analysis. **A:** Histogram of unadjusted p-values from single gene analysis of islet autoimmunity positive vs. age-matched children who did not develop islet autoimmunity. **B:** Histogram of p-values after multiple testing, controlling the false discovery rate. **C:** Sorted −log10(p-values) of pathway enrichment for islet autoimmunity status. Dashed line indicates the multiple testing threshold at an FDR of 0.1. Abbreviations: Ab+ = Islet autoantibody-positive, Ab− = Islet autoantibody negative.

Pathway enrichment analysis on KEGG pathways identified 20 as significantly regulated (FDR <= 0.1, Figure 3C and Supplemental file 2). The top-ranked pathways were “p53 signaling pathway”, “Ubiquitin mediated proteolysis”, and “RIG-I-like receptor signaling pathway.” The two significant pathways from the Cesarean section analysis, “Pyrimidine metabolism” and “Pentose phosphate pathway”, also appeared as significant in the islet autoantibody analysis. Notably, comparing gene expression samples of children before seroconversion (up to 6 months before) and age-matched children without islet autoantibodies yielded no significant genes or pathways (Supplemental file 3). In addition, we investigated children that had gene expression samples both before and after seroconversion (up to 6 month before and after) in a paired analysis, leaving only seven children. No genes or pathways were found to be differentially expressed (Supplemental file 3).

Taken together, we observed several immune system-related and nucleotide synthesis pathways associated with islet autoimmunity, which included the pathways found in our Cesarean section analysis.

### Coherent gene expression changes between Cesarean section and islet autoimmunity

We further investigated the similarities of transcriptional changes between the two risk factors. First, we found that the individual gene regulation of both “Pyrimidine metabolism” and the “Pentose phosphate pathway” pathway showed similar patterns of up- and down-regulation between Cesarean section and islet autoimmunity (Figure 4A, Supplemental file 4).

**Figure 4:**
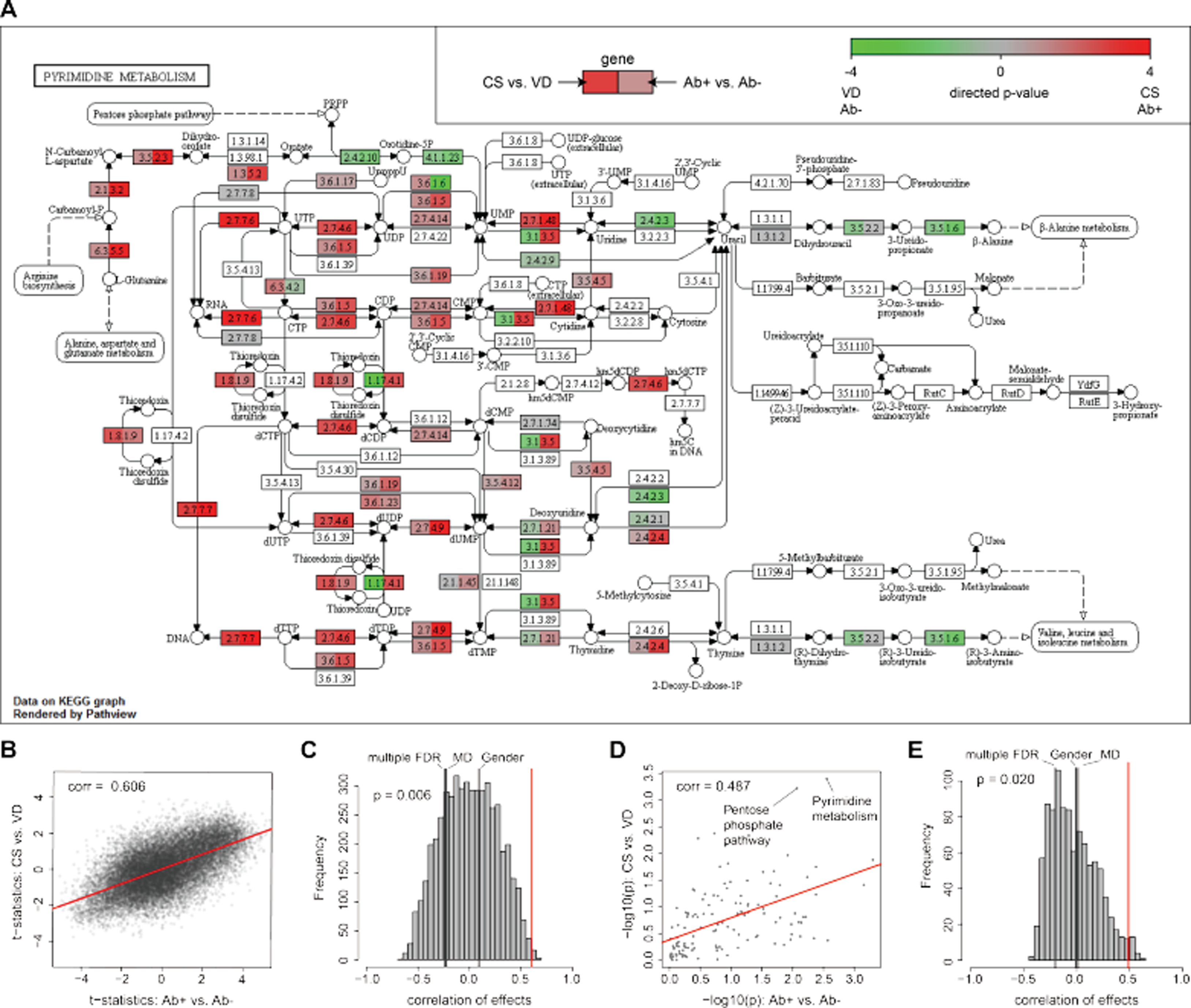
Comparison of Cesarean section and islet autoimmunity signatures. **A:** The pyrimidine metabolism pathway is shown as an example, with two-sided node coloring according to directed p-values (−log10(p) * sign(t-statistic)). **B:** T-statistics per gene from both analyses; the x-axis indicates results from islet autoimmunity status and the y-axis the results from Cesarean section vs. vaginal delivery. **C:** Empirical distribution of correlation coefficients between Cesarean section and permuted class labels of islet autoimmunity status for 5,000 permutations. The red line indicates the ‘true’ correlation between the results from the analysis of Cesarean section and islet autoimmunity status. Black lines indicate the correlation between Cesarean section and multiple first-degree relatives (multiple FDR), maternal diabetes (MD), and gender. **D:** Correlation of pathway directed p-values in Cesarean section analysis and islet autoimmunity status analysis. **E:** Empirical distribution of correlation coefficients between Cesarean section and permuted class labels of islet autoimmunity status at pathway level for 1,000 permutations. The red line indicates the ‘true’ correlation between Cesarean section pathways and islet autoimmunity status pathways. Abbreviations: CS = Cesarean section, VD = Vaginal delivery, Ab+ = Islet autoantibody-positive, Ab− = Islet autoantibody negative.

Extending this analysis, we quantified the relationship of gene regulation between the two factors at a systematic level. We correlated the standardized effects from Cesarean section and development of islet autoantibodies across all genes (Figure 4B), revealing a striking correlation of 0.606 (p = 0.0062, Figure 4C). In other words, genes regulated by Cesarean section, despite the rather weak signal strength in our study, were also regulated in the same direction by the initiation of islet autoimmunity. A similar correlation between the effects of Cesarean section and islet autoimmunity was observed at the pathway level, with a correlation of 0.49 (p = 0.02, Figure 4D+E), supporting the functional agreement of transcriptional changes between the two risk factors. Importantly, we did not observe that children born by Cesarean section developed islet autoantibodies more frequently than children born by vaginal delivery, ruling out a confounding effect not related to gene expression (Supplemental file 1: Table S1).

In contrast to the profound Cesarean section to islet autoantibodies correlation at transcript level, we found the effects of gender, maternal diabetes and multiple first-degree relatives to be randomly correlated with Cesarean section at the single gene level (maternal diabetes, r = −0.22, p = 0.53; gender, r = 0.09, p = 0.78; multiple first-degree relatives, r = −0.24, p = 0.49) and at the pathway level (maternal diabetes, r = 0.01, p = 0.97; gender, r = −0.01, p = 0.97; multiple first-degree relatives, r = −0.23, p = 0.34), as shown in Figure 4 C+E and Supplemental file 5. This demonstrates the specificity of the Cesarean section–islet autoantibody effect correlation.

In summary, this analysis showed that the gene expression changes in these two risk factors, Cesarean section and islet autoimmunity, are remarkably coherent.

### Signatures of immune cell activation

The pentose phosphate pathway is a universal, central metabolic pathway in the cytosol, which supports cell proliferation and survival [15]. The non-oxidative branch of the pentose phosphate pathway branches off glycolysis and generates ribose 5-phosphate as a precursor for the synthesis of nucleotides and amino acids necessary for cell growth and division. Moreover, the pyrimidine metabolism pathway is directly related to the pentose phosphate pathway in its role in nucleotide synthesis. Since we investigated PMBCs, these differentially regulated pathways in Cesarean section and islet autoimmunity point toward a general activation of immune cells. A direct proof of this hypothesis in the same children was unfeasible in the context of this study. Instead, we collected evidence from several secondary analysis steps.

First, we investigated whether regulated genes from both analyses enriched immune genes annotated in innateDB [16]. Indeed, there was a significant accumulation of higher standardized effects for immune genes compared to non-immune genes, for both Cesarean section (Wilcoxon rank sum test: p = 0.0002) and autoimmunity (p = 2.6 × 10^−15^); see Supplemental file 6.

Second, we compared results from the mixture of PBMC cells with published data on activation in isolated immune cells. For the analysis, we used transcriptomics data from isolated naïve and activated human CD4+ T cells [17]. We calculated gene expression differences before and after activation, and applied enrichment analysis to identify differentially expressed pathways. In particular, the “Pentose phosphate pathway” (p = 0.049) and “Pyrimidine metabolism” (p = 0.035) pathways were significantly differentially expressed in activated CD4+ T cells (Supplemental file 2). To compare the effects of CD4+ T cell activation with effects from the Cesarean section and islet autoantibody analyses, we calculated correlations of the standardized effects. We observed borderline significant correlations between changes in activated CD4+ T cell and Cesarean section (r = 0.20, p = 0.049, Figure 5 A+B) and islet autoimmunity (r = 0.24, p = 0.052, Figure 5 C+D). Remarkably, the association was substantially stronger at pathway level for both Cesarean section (r = 0.45, p = 0.021, Figure 5 E+F) and islet autoimmunity (r = 0.57, p = 0.008, Figure 5 G+H). The pathway association was replicated in a second transcriptomics dataset from naïve and activated CD4+ T cells, monocytes, and natural killer cells, published in [18]. Detailed results are shown in Supplemental file 7.

**Figure 5:**
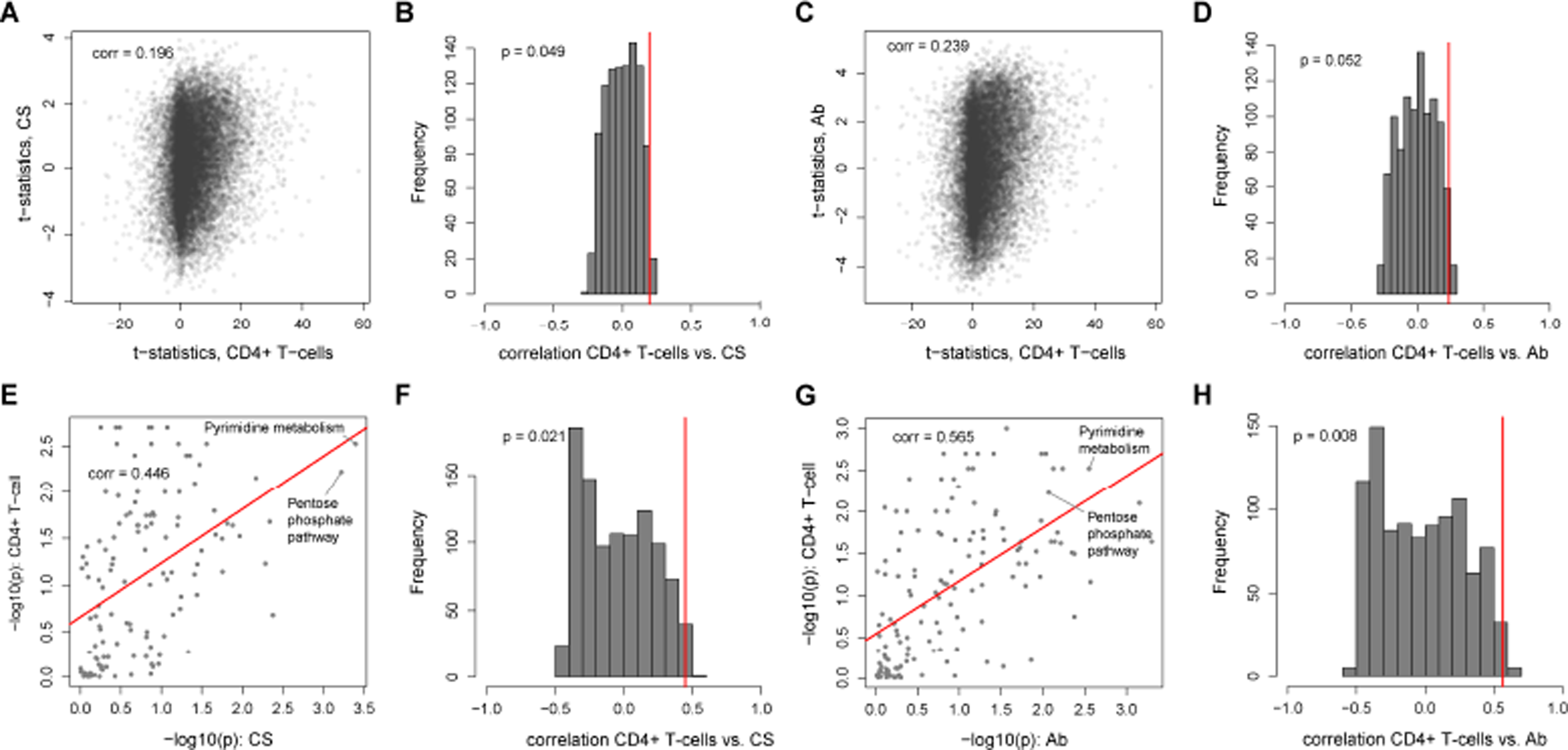
Signatures of immune cell activation-. **A:** Correlation between single gene effects in Cesarean section (CS) compared to the association between naïve and activated CD4+ cells. **B:** Histogram of correlation of permuted class labels of CD4 T cells and Cesarean section at the single gene level and the “true” correlation effect. **C:** Correlation between single gene effects in islet autoimmunity status (Ab) compared to the association between naïve and activated CD4+ cells. **D:** Histogram of correlation of permuted class labels of CD4 T cells and islet autoimmunity status at the single gene level and the “true” effect. **E:** Correlation between pathway effects in Cesarean section compared to the association between naïve and activated pathways for CD4+ cells. **F:** Histogram of correlation of permuted class labels of CD4 T cells and Cesarean section at the pathway level and the “true” effect. **G:** Correlation between pathway effects in islet autoimmunity status compared to the association between naïve and activated pathways for CD4+ cells. **H:** Histogram of correlation of permuted class labels of CD4 T cells and islet autoimmunity status at the pathway level and the “true” effect.

In summary, we observed a significant correlation between the functional effects of human lymphocyte activation and the effects of Cesarean section and islet autoimmunity on gene expression in PBMCs.

## Discussion

We identified coherent gene expression signatures for Cesarean section, an early risk factor for type 1 diabetes, and islet autoantibody positivity, an obligatory stage of autoimmune response prior to the development of autoimmune type 1 diabetes. Specifically, at the transcriptome level, we identified two pathways involved in nucleotide synthesis and cell proliferation that were regulated in PBMCs of children born by Cesarean section. This analysis required an extended statistical model to incorporate the complex time information in our present dataset. Islet autoantibody analysis revealed various pathways generally involved in immune processes, including the aforementioned nucleotide synthesis pathways. Comparing global gene expression signatures, we found that transcriptomic changes were systematically and significantly correlated between the Cesarean section and islet autoantibody-positive analyses. Importantly, transcriptional signatures of Cesarean section did not correlate with gender, maternal diabetes, or multiple first-degree relatives, demonstrating that the correlation is specific to Cesarean section and islet autoimmunity. In a functional follow-up analysis, both Cesarean section and islet autoimmunity signatures were significantly correlated with gene expression changes observed during activation of isolated CD4+ T lymphocytes. At pathway level, this correlation was also observed for monocytes and natural killer cells in a second dataset.

We can speculate on the biological basis of our statistical observations. The coherent regulation of proliferation pathways in blood PBMCs may indicate a general activation of the immune system and immune cells for both Cesarean section and seroconversion. In the case of Cesarean section, this activation might indirectly reflect different microbial exposures during birth. This idea is indirectly supported by findings that the microbiome is affected by the mode of delivery, and in turn affects the developing immune system [9, 10]. For the autoimmunity results, the precise interplay and timing of the occurrence of environmental stimuli, immune response and the development of islet autoantibodies still need to be elucidated. Regarding a hypothetical disease trajectory leading to type 1 diabetes, it is conceivable that the observed gene expression signatures may reflect a transient deflection of the immune system and that this primes a child for subsequent progression to type 1 diabetes in the presence of other risk factors.

An interesting general observation in our analysis was the increased correlation of gene expression signatures when performing pathway analysis instead of single gene analysis. This pattern was observed both for the comparison of Cesarean section and islet autoantibody-positive signatures, and for comparisons of these two signatures with the isolated immune cells. These findings indicate that pathway analysis substantially reduced the noise compared to single gene analysis, which allowed us to identify common patterns at a functional level.

Our study could be extended and improved in several directions. (1) The present dataset has rather low statistical power for the islet autoantibody analysis, with only 15 positive samples available for at most 6 months post-seroconversion. While differential expression changes were remarkably significant, the analysis should be validated with a larger sample size. (2) The hypothesis of coherent immune system activation should ideally be confirmed in an isolated primary T cell population from children after Cesarean section or after seroconversion. We took an indirect route using PBMCs rather than a more general activation experiment with isolated immune cells. (3) The incorporation of further environmental factors, such as nutrition and medical parameters, in combination with the children’s genetic background, is expected to give a more complete picture of the parameters leading to autoimmunity and type 1 diabetes.

## Conclusions

In summary, we found a transcriptional link between Cesarean section and islet autoimmunity, pointing toward a transiently altered immune system in the susceptible period of islet autoimmunity generated by Cesarean section, which was remarkably coherent with the changes observed after islet autoimmunity.

## Methods

### PBMC gene expression data

We used the BABYDIET PBMC gene expression data deposited in ArrayExpress (accession number: E-MTAB-1724) [6], in combination with non-public data on Cesarean section, islet autoimmunity, family history, gender and age. In this data set, 454 samples from 109 children were available. One individual of the original 109 children was removed in the Cesarean section analysis, since no information about the type of delivery was recorded. Raw gene expression data of 33,297 probes were normalized using the Robust Multi-Array Average (RMA) method [19]. Probes without annotation in the Affymetrix hugene11 data and duplicates were removed using the R package *genefilter*, leaving 18,720 genes for further analysis. Age at sampling ranged from 0.21 years to 9.15 years, with a median of 1.53 years.

### Generalized additive mixed model for time-resolved data

Transcriptomics samples were available for multiple time points per child. To model gene expression from multiple, non-matching timepoints in relation to Cesarean section in a joint approach, we employed a generalized additive mixed model (GAMM) [20]. In this GAMM, the model structure is defined as

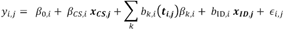

where *y_i,j_* is the gene expression of sample *j* for gene *i, β_0,i_* is the intercept or gene-wise average expression, *β_cs,i_* describes the effect of Cesarean section on gene expression with *x_cs,j_* = 1 for Cesarean section and *x_cs,j_* = 0 for vaginal delivery. Adjusting for the multiple measurements in time *t_i,j_*, a spline function was included as ∑_*k*_*b_k,i_*(***t_i,j_***)*β_k,i_* per gene over all samples, where *k* is the estimated number of basis functions, *b_k,i_* are the basis functions of the spline and *β_k,i_* their regression coefficients. In addition, we included a random effect as *b_ID,i_*~ *N*(0,*σ_i_*), with *σ_i_* estimated per gene, correcting for the dependence of several measurements per child, denoted as ***x_id,j_***, and an error term *ϵ_i,j_* as i.i.d. normally distributed noise. The noise *ϵ_i,j_* and the noise of the subject specific random effect *σ_i_* were assumed to be independent. Inference of the GAMM was performed using a restricted maximum likelihood approach [21] using the R package *mgcv*.

### Linear model for seroconversion analysis

To investigate differences in gene expression between children after development of islet autoantibodies and age-matched children who did not develop islet autoantibodies, we selected the first sample up to six months after development of islet autoimmunity for each child, when such a sample was available (see Figure 1B). These children were compared to all available age-matched children who did not develop autoantibodies, by applying a linear regression model per gene:

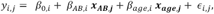

where *y_i,j_* is the gene expression of child *j* for gene *i*, *β_0,i_* the intercept, *β_AB,i_* describes the effect on gene expression of islet autoimmunity, with ***x_AB,j_*** = 1 for islet autoantibody-positives and ***x_AB,j_*** = 0 for islet autoantibody-negatives, and ***β_age,i_*** the effect of age ***x_age,j_*** on gene expression. The residual term *ϵ_i,j_* was defined as i.i.d. normally distributed noise.

### Pathway enrichment analysis

We performed pathway enrichment analysis based on the KEGG database [12]. We identified differentially regulated pathways using the “Significance Analysis of Function and Expression” (SAFE) algorithm [22], R package *safe*. In this approach, p-values from local gene tests are computed and Wilcoxon rank sum statistics assess whether local statistics are systematically increased in the pathway, compared to the background (global test). Local gene tests were calculated on the residuals of models from equations (1) and (2), but excluding the term for the factor of interest (Cesarean section or islet autoimmunity). SAFE uses a sample permutation-based approach for p-value calculation, which avoids false positive results due to correlating transcripts. The number of permutations was set to 5,000.

For node coloring of pathways (Figure 4A+B), we used a “directed p-value”, defined as (−log10(p-value) * sign(regression coefficient)). For nodes in the pathway that had multiple genes annotated, the highest effect was shown.

### Comparing Cesarean section and islet autoimmunity results

To compare the results obtained from the Cesarean section and islet autoimmunity (Ab+) analyses, we calculated the Pearson correlation between the standardized effect estimates of both analyses. Standardized effect estimates were represented by the t-statistic in the single gene analysis, and a “directed p-value” (analogously to previous section) in the pathway analysis. To assess the statistical significance of the correlation, we calculated

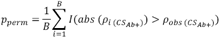

with B being the number of permutations, *ρ*_*obs* (*CS*_*Ab*+_)_ the observed “true” correlation of effects between both analyses, *I()* the indicator function and *ρ*_*i* (*CS*_*Ab*+_)_ the correlation between the effects of Cesarean section and permuted islet autoimmunity status. The number of permutations was fixed at B = 5,000 for the single gene analysis and 1,000 for the pathway analyses. The same analysis with 1,000 permutations was repeated for the factors maternal diabetes, gender, and multiple first-degree relatives.

### Immune cell activation analysis

To investigate the enrichment of genes within annotated immune genes from innateDB [16], we calculated Wilcoxon rank sum tests using the t-statistics from the single gene level analysis (see above) to identify differences between the distribution of immune genes and non-immune genes. For comparison of standardized effects, we used two published datasets, one from isolated CD4+ T cells before and after activation (GEOD-33272, processed data downloaded) and one containing different subsets of human leukocytes before and after activation (GEOD-22886, processed data downloaded). Single gene differential analysis was performed on log2-transformed expression values using linear regression, with gene expression as the response, and activation status as the explaining variables. For SAFE pathway enrichment, the number of permutations was set to 1,000. To calculate permutation-based p-values of Pearson correlations, we used 1,000 permutations.

All statistical analyses were performed using the computing environment R version 3.3.2 [23].

## List of abbreviations

PBMC: peripheral blood mononuclear cells
GAMM: generalized linear mixed model

## Acknowledgments

We thank Dr. Nikola Müller and Dr. Steffen Sass for sharing their expertise on transcriptomics data.

## Declarations

### Ethics approval and consent to participate

The BABYDIET study was approved by the ethics committee of Ludwig-Maximilians University (Protocol No. 329/00) and is registered at ClinicalTrials.gov NCT01115621. The parents or guardians of each child provided informed consent for participation in the BABYDIET study. All samples and information were collected after obtaining signed informed consent.

### Consent for publication

Not applicable.

### Availability of data and materials

This publication uses publicly available gene expression datasets, available via ArrayExpress (accession number E-MTAB-1724) and the Gene Expression Omnibus (accession numbers GSE33272 and GSE22886). The informed consent given by study participants does not cover data posting of the clinical covariates in public databases.

## Authors’ contributions

Study concept and design: ML, AGZ, JK. Statistical analysis: ML. Acquisition of data: RP, RF, CW, AGZ. Analysis and interpretation of data: ML, EB, JK. Drafting of the manuscript: ML, EB, AGZ, JK. Critical revision of the manuscript for important intellectual content: ML, RP, RF, FJT, CW, EB, AGZ, JK. All authors read and approved the final manuscript.

## Supplemental files

Supplemental file 1: Detailed information on longitudinal sampling and on Cesarean section in the dataset.

Supplemental file 2: Statistical results of all gene and pathway analyses.

Supplemental file 3: Results of analysis for samples up to 6 months before islet autoimmunity compared to age matched islet autoantibody negatives children. Results of paired analysis of samples before and after seroconversion.

Supplemental file 4: Pentose phosphate pathway with two-sided node colouring according to directed p-values.

Supplemental file 5: Permutation results for maternal diabetes, gender and multiple first-degree relative.

Supplemental file 6: Gene list downloaded from innateDB filtered for human genes. T-statistics of annotated and not annotated genes are shown in a violin plot.

Supplemental file 7: Detailed single gene and pathway results of transcriptomics dataset GEO-22886 of activated immune cells

## References

1. Donath MY, Halban PA: Decreased beta-cell mass in diabetes: significance, mechanisms and therapeutic implications. Diabetologia 2004, 47(3):581-589.

2. Krischer JP, Lynch KF, Schatz DA, Ilonen J, Lernmark A, Hagopian WA, Rewers MJ, She JX, Simell OG, Toppari J et al: The 6 year incidence of diabetes-associated autoantibodies in genetically at-risk children: the TEDDY study. Diabetologia 2015, 58(5):980-987.

3. Giannopoulou EZ, Winkler C, Chmiel R, Matzke C, Scholz M, Beyerlein A, Achenbach P, Bonifacio E, Ziegler AG: Islet autoantibody phenotypes and incidence in children at increased risk for type 1 diabetes. Diabetologia 2015, 58(10):2317-2323.

4. Lipman TH, Levitt Katz LE, Ratcliffe SJ, Murphy KM, Aguilar A, Rezvani I, Howe CJ, Fadia S, Suarez E: Increasing incidence of type 1 diabetes in youth: twenty years of the Philadelphia Pediatric Diabetes Registry. Diabetes care 2013, 36(6):1597-1603.

5. Kallionpaa H, Elo LL, Laajala E, Mykkanen J, Ricano-Ponce I, Vaarma M, Laajala TD, Hyoty H, Ilonen J, Veijola R et al: Innate immune activity is detected prior to seroconversion in children with HLA-conferred type 1 diabetes susceptibility. Diabetes 2014, 63(7):2402-2414.

6. Ferreira RC, Guo H, Coulson RM, Smyth DJ, Pekalski ML, Burren OS, Cutler AJ, Doecke JD, Flint S, McKinney EF et al: A type I interferon transcriptional signature precedes autoimmunity in children genetically at risk for type 1 diabetes. Diabetes 2014, 63(7):2538-2550.

7. Cardwell CR, Stene LC, Joner G, Cinek O, Svensson J, Goldacre MJ, Parslow RC, Pozzilli P, Brigis G, Stoyanov D et al: Caesarean section is associated with an increased risk of childhood-onset type 1 diabetes mellitus: a meta-analysis of observational studies. Diabetologia 2008, 51(5):726-735.

8. Bonifacio E, Warncke K, Winkler C, Wallner M, Ziegler AG: Cesarean section and interferon-induced helicase gene polymorphisms combine to increase childhood type 1 diabetes risk. Diabetes 2011, 60(12):3300-3306.

9. Biasucci G, Benenati B, Morelli L, Bessi E, Boehm G: Cesarean delivery may affect the early biodiversity of intestinal bacteria. The Journal of nutrition 2008, 138(9):1796S-1800S.

10. Caicedo RA, Schanler RJ, Li N, Neu J: The developing intestinal ecosystem: implications for the neonate. Pediatric research 2005, 58(4):625-628.

11. Neu J, Rushing J: Cesarean versus vaginal delivery: long-term infant outcomes and the hygiene hypothesis. Clinics in perinatology 2011, 38(2):321-331.

12. Kanehisa M, Goto S: KEGG: kyoto encyclopedia of genes and genomes. Nucleic acids research 2000, 28(1):27-30.

13. Jiang P, Du W, Wu M: Regulation of the pentose phosphate pathway in cancer. Protein & cell 2014, 5(8):592-602.

14. Lane AN, Fan TW: Regulation of mammalian nucleotide metabolism and biosynthesis. Nucleic acids research 2015, 43(4):2466-2485.

15. O’Neill LA, Kishton RJ, Rathmell J: A guide to immunometabolism for immunologists. Nature reviews Immunology 2016, 16(9):553-565.

16. Breuer K, Foroushani AK, Laird MR, Chen C, Sribnaia A, Lo R, Winsor GL, Hancock RE, Brinkman FS, Lynn DJ: InnateDB: systems biology of innate immunity and beyond--recent updates and continuing curation. Nucleic acids research 2013, 41(Database issue):D1228-1233.

17. Maliga Z, Junqueira M, Toyoda Y, Ettinger A, Mora-Bermudez F, Klemm RW, Vasilj A, Guhr E, Ibarlucea-Benitez I, Poser I et al: A genomic toolkit to investigate kinesin and myosin motor function in cells. Nature cell biology 2013, 15(3):325-334.

18. Abbas AR, Baldwin D, Ma Y, Ouyang W, Gurney A, Martin F, Fong S, van Lookeren Campagne M, Godowski P, Williams PM et al: Immune response in silico (IRIS): immune-specific genes identified from a compendium of microarray expression data. Genes and immunity 2005, 6(4):319-331.

19. Irizarry RA, Hobbs B, Collin F, Beazer-Barclay YD, Antonellis KJ, Scherf U, Speed TP: Exploration, normalization, and summaries of high density oligonucleotide array probe level data. Biostatistics 2003, 4(2):249-264.

20. Wood SN: Generalized additive models: an introduction with R: CRC press; 2017.

21. Wood SN: Fast stable restricted maximum likelihood and marginal likelihood estimation of semiparametric generalized linear models. Journal of the Royal Statistical Society: Series B (Statistical Methodology) 2011, 73(1):3-36.

22. Barry WT, Nobel AB, Wright FA: Significance analysis of functional categories in gene expression studies: a structured permutation approach. Bioinformatics 2005, 21(9):1943-1949.

23. Team RC: R: a language and environment for statistical computing. R Development Core Team, Vienna. In.; 2016.

